# Transcriptomic signatures and network-based methods uncover new Senescent Cell Anti-Apoptotic Pathways and Senolytics

**DOI:** 10.1101/2024.05.28.596326

**Authors:** Samael Olascoaga, Mina Konigsberg, Jesús Espinal-Enríquez, Hugo Tovar, Félix Matadamas-Martínez, Jaime Pérez-Villanueva, Norma Edith López-Diazguerrero

## Abstract

Cellular senescence is an irreversible cell cycle arrest caused by various stressors that damage cells. Over time, senescent cells accumulate and contribute to the progression of multiple age-related degenerative diseases. It is believed that these cells accumulate partly due to their ability to evade programmed cell death through the development and activation of survival and anti-apoptotic resistance mechanisms; however, many aspects of how these survival mechanisms develop and activate are still unknown. By analyzing transcriptomic signature profiles generated by the LINCS L1000 project and using network-based methods, we identified various genes that could represent new senescence-related survival mechanisms. Additionally, employing the same methodology, we identified over 600 molecules with potential senolytic activity. Experimental validation of our computational findings confirmed the senolytic activity of Fluorouracil, whose activity would be mediated by a multi-target mechanism, revealing that its targets AURKA, EGFR, IRS1, SMAD4, and KRAS are new senescence-associated survival and anti-apoptotic resistance pathways. The development of these pathways could depend on the stimulus that induces cellular senescence. The SCAPs development and activation mechanisms proposed in this work offer new insights into how senescent cells survive. Identifying new anti-apoptotic resistance targets and drugs with potential senolytic activity paves the way for developing new pharmacological therapies to eliminate senescent cells selectively.

**Graphical Abstract:** 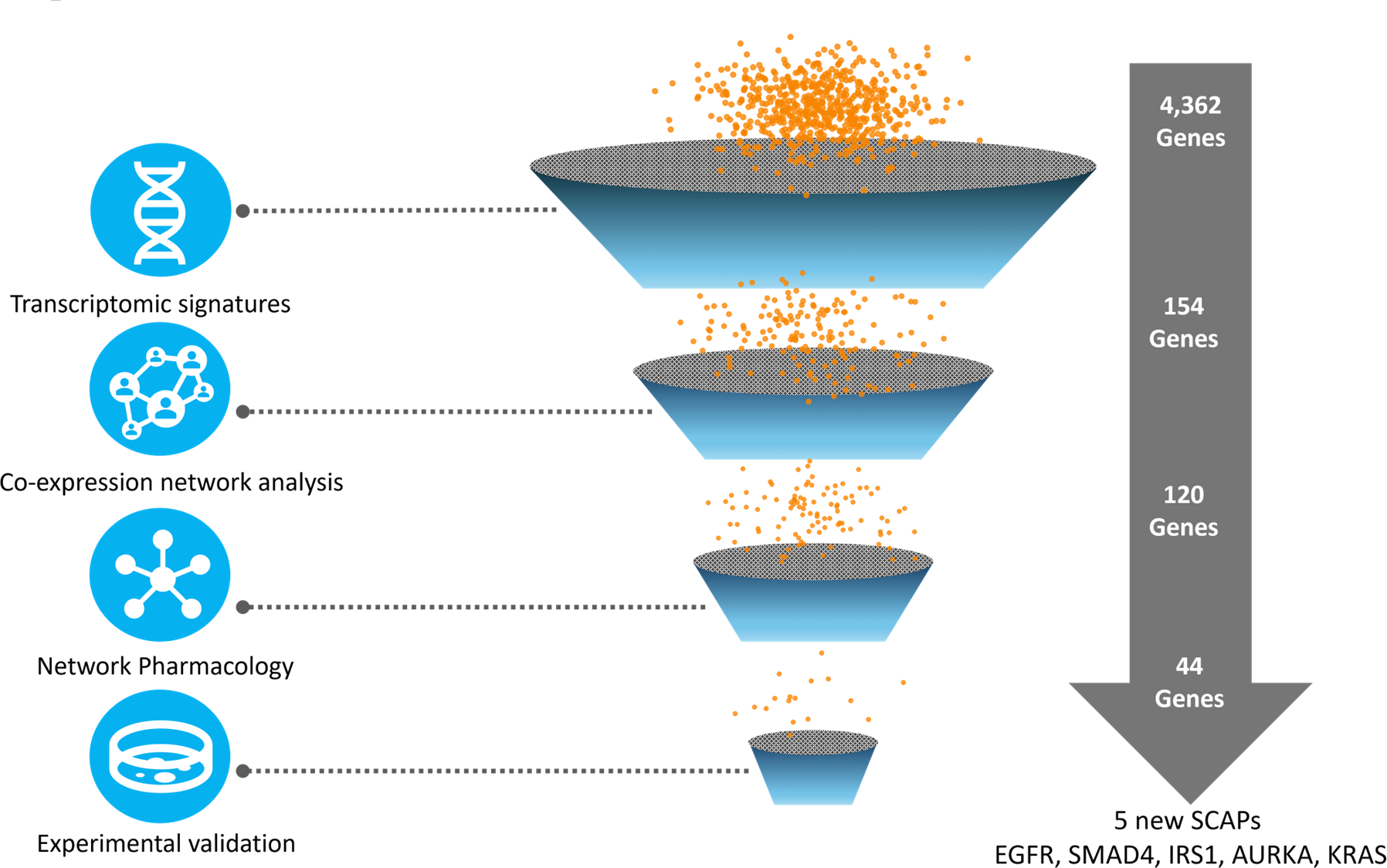

## Introduction

Cellular senescence (CS) is a state of irreversible cell cycle arrest in response to various stressors that can damage cells. These agents include telomere shortening, oncogenic activation, ionizing radiation, and oxidative stress (Herranz & Gil, 2018). Senescent cells (SCs) exhibit various morphological and molecular alterations, such as acquiring an elongated and flattened phenotype in cell cultures, accumulating lysosomes and misfolded proteins in the cytoplasm, and alterations in mitochondrial dynamics. Additionally, they also secrete various soluble components into the extracellular medium, such as cytokines, chemokines, growth factors, and metalloproteins, among others, collectively known as the senescence-associated secretory phenotype (SASP), which can profoundly modify and influence the cellular microenvironment (Childs et al., 2015). CS is widely recognized as a tumor suppressor mechanism (Coppé et al., 2010); however, it has also been demonstrated that the accumulation of SCs is implicated in the onset and progression of multiple chronic-degenerative diseases associated with aging, including cancer (Wissler Gerdes et al., 2020). Senescent cells cannot respond to pro-apoptotic stimuli, leading to their accumulation in organs and tissues over time (Soto-Gamez et al., 2019) due to the development and activation of various survival and anti-apoptotic resistance pathways (SCAPs) (Herranz & Gil, 2018), such as the PI3K/AKT, p53, BCL-2 family, and heat shock proteins (HSPs) signaling pathways, among others (Hu et al., 2022). The first SCAPs were discovered through bioinformatic analysis of transcriptomic data generated from gene inhibition with small interfering RNAs (siRNAs). In total, 39 genes were found that induced selective death of SCs when individually inhibited. This finding led to the discovery of the first senolytic molecules, Dasatinib and Quercetin, which are small molecules capable of inducing selective death of SCs through various mechanisms (Zhu et al., 2015).

The Library of Integrated Network-based Cellular Signatures (LINCS L1000) project is a biomedical initiative focused on generating molecular signatures of human cells in response to various chemical and genetic perturbations. It utilizes a large-scale gene expression technology to measure changes in the expression of approximately 1,000 highly representative genes. These genes have been carefully selected to represent the global human transcriptome. The idea is that analyzing the changes in these genes after exposure to different drugs or genetic modifications makes it possible to infer broader patterns of gene activity (Subramanian et al., 2017).

In this work, our objective was to identify new SCAPs. For this purpose, we analyzed the transcriptomic signatures generated by the LINCS L1000 project, where individual genes are silenced using short hairpin RNAs (shRNAs). These transcriptomic profiles provide a rich and detailed database to analyze gene expression changes associated with the inhibition of genes that, when inhibited, generate the same biological response. Thus, we pinpointed genes whose suppression elicits a transcriptomic response akin to that observed when known SCAPs are inhibited. Then, with these genes, we constructed and analyzed gene co-expression networks (GCNs) using gene expression profiles from human lung samples to determine if the newly identified genes are functionally related to each other and known SCAPs, which would place the identified genes as probable SCAPs. We then used network pharmacology to identify inhibitors of the proteins encoded by the new SCAPs candidates. The computational findings were experimentally validated using two distinct models of senescence in primary cultures of normal human lung fibroblasts, one of replicative senescence (RS) and the other of stress-induced premature senescence (SIPS). Two of the drugs identified through network-based methods inhibited the new SCAPs in the two senescence models. Figure 1 graphically summarizes the general strategy used in this work.

**Figure 1.**
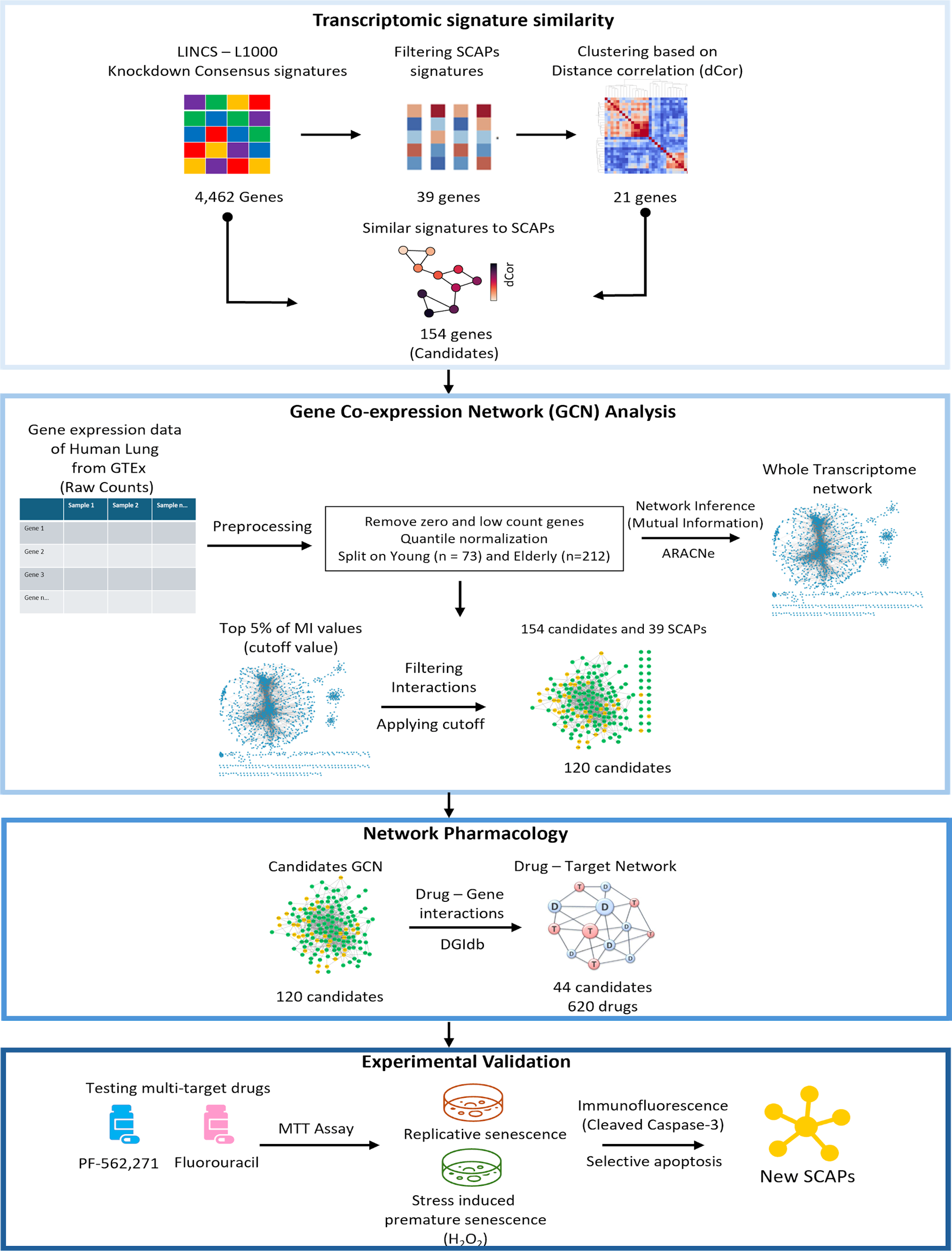
General Methodology. Network-based computational approaches were combined with experimental methods for the identification of new SCAPs.

## Methods

### Similarity of Transcriptomic Signatures

Consensus transcriptomic signatures data were used, condensing the signature profiles generated by the individual shRNA silencing of 4,462genes in 9 different cell lines (Daniel Himmelstein, n.d.). Of the original 39 SCAPs, 29 were available within the signature’s dataset. Only the 978 landmark genes quantified in the original LINCS L1000 platform were considered for the similarity calculations.

The distance correlation (dCor) was chosen as a similarity measure, which considers the difference between the joint and marginal distributions of two variables. Unlike Pearson’s correlation, which only measures linear relationships, dCor can detect nonlinear and more complex interactions. A zero value indicates total independence, while higher values indicate greater dependence (Székely et al., 2007). The dCor was calculated using the distance correlation function implemented in the dCor 0.5 package (Ramos-Carreño & Torrecilla, 2023), and hierarchical clustering analysis was performed using the clustermap function and the average method implemented in the seaborn 0.11.2 package (Waskom, 2021).

The statistical significance of the similarity of the SCAPs signatures was obtained by bootstrapping (Henderson, 2005). The dCor of each pair of genes within the original set of 4,462signatures was calculated, then random samples of the same size as the reference sample were taken, and the number of times the random samples had a median dCor higher than the reference was calculated through 100,000 simulations.

We randomly selected 22 genes (simulating 21 SCAPs and 1 candidate) and calculated their median dCor values. After one million simulations, we used the highest random dCor value (0.405) as the cutoff. The visualization of the genes whose signature turned out to be similar to that of the SCAPs was carried out through a network generated using Cytoscape 3.10.1 (Shannon et al., 2003).

### Data Acquisition

The Genotype-Tissue Expression (GTEx) project provides gene expression data generated through bulk RNA-seq from 53 different tissues, collected from 714 donors without reported diseases whose ages ranged from 20 to 79 years (Aguet et al., 2020). Specifically, from the GTEx V8 dataset, 578 human lung samples from men and women were obtained. Each sample included raw counts for a total of 56,200 genes.

### Data Preprocessing

The raw gene expression count values from human lung samples were preprocessed as follows: Initially, the mean of the counts for each gene across the samples was calculated. Genes with a mean below 10 were removed, and genes with zero count values in more than 50% of the samples were discarded (García-Cortés et al., 2022). Subsequently, the dataset was quantile normalized using the qnorm 0.8.1 library. The next step consisted of dividing the dataset into two groups according to the ages defined by the GTEx project: young (20 to 39 years old, n = 73) and old (60 - 70 years old, n = 212). Protein-coding genes, long non-coding RNAs, microRNAs, pseudogenes, and other types of RNA species were retained. After this preprocessing, each sample retained 21,096 genes.

### Gene Co-expression Network (GCN) Inference

GCNs were inferred using mutual information (MI) to measure gene co-expression. Mutual information is a statistical measure that quantifies how much information one random variable, such as gene expression, contains about another. The ARACNe (Algorithm for the Reconstruction of Gene Regulatory Networks) algorithm, one of the most popular methods for inferring GCNs, calculates the MI between two data series (Margolin et al., 2006)). ARACNe was applied to both preprocessed datasets (young and old) to establish correlations between gene pairs. The complete transcriptome network was calculated for both datasets, spanning 21,096× 21,096 gene pairs. To speed up the calculation, we used the multi-core C++ version without adaptive partitioning inference (Andonegui-Elguera et al., 2021). This version is available at https://github.com/josemaz/aracne-multicore

The MI value corresponding to the 95th percentile of each network (young and old) was obtained using the MI values of the complete transcriptome. These values were then used as cutoff points in each GCN constructed in this work to get statistically significant networks.

### Network Pharmacology and Drug Repositioning

To construct the network of senolytic drugs and their targets, 59 senolytic drugs and 550 targets reported in previous work were used (Olascoaga-Del Angel et al., 2022). We applied bootstrapping to determine whether the genes from the GCN were enriched in senolytic drug targets. Specifically, the 4,462 genes from the LINCS L1000 dataset were used as background, and one million simulations were performed. In each simulation, 120 elements were randomly selected (the same number as the genes in the GCN without considering the original SCAPs genes), and the frequency with which these random elements matched elements within the network of 550 senolytic drug targets was calculated.

The Drug Gene Interaction Database (DGIdb) v4.2.0 (Freshour et al., 2021) was used for drug repositioning. This database compiles information from 22 other pharmacological databases (consulted on 03/05/2024). The search for molecules for the 120 GCN genes resulted in 620 drugs interacting uniquely with 44 of the 120 GCN genes.

### Cell Culture

Human lung fibroblasts from the CCD-8Lu primary cell line, acquired from ATCC (catalog number: CCL-201), were cultured in 100 mm Petri dishes. The cells were maintained in Gibco™ DMEM/F-12 culture medium (catalog number: 11320033), supplemented with 10% heat-inactivated fetal bovine serum (FBS) from Gibco™ (catalog number: 16000044) and 1% penicillin, amphotericin B, and streptomycin, also from Gibco™ (catalog number: 15240-062). The cells were incubated at 37°C with 5% CO_2_.

### Stress-Induced Premature Senescence (SIPS)

Approximately 10,000 cells with PD < 10 were seeded in 48-well plates and incubated for 2 hours. After this time, most cells adhered to the culture plate and were subsequently exposed to H_2_O_2_ (Sigma-Aldrich, cat. H1009) at 100, 150, and 200 μM concentrations. After 2 hours, two washes with 1X PBS were performed, and fresh medium was added. At 24 hours, the medium was renewed. The cells were incubated at 37°C with 5% CO_2_. On subsequent days (3, 6, and 9) after H_2_O_2_ treatment, the cells were detached using 0.05% trypsin. Cell viability was evaluated in three independent experiments using the trypan blue technique, each in triplicate.

Statistical analysis was performed with *Prism 8.0.2*. The Shapiro-Wilk test was applied for normality analysis, with a significance level of *p* < 0.05. A two-way ANOVA was used to analyze the peroxide concentration that reduces proliferation over 9 days, followed by a Tukey post-hoc test, using a significance level of *p* < 0.05.

### Replicative Senescence (RS)

To establish a standardized replicative senescence (RS) model, cells at a low passage number (<10) were seeded into 6-well plates, and the culture medium was refreshed every 48 hours. Once the cells attained approximately 80% confluency, they were detached using 500 μL of 0.05% trypsin per well. Subsequently, the cells were reseeded at a 1:4 ratio across new 6-well plates, a subculturing process where cells from the original culture were divided into four separate plates. This subculturing process was repeated until the cells achieved a high passage number (> 60). Upon reaching passage 60, 10,000 high-passage cells were seeded in new plates and allowed to stabilize for 24 hours to accommodate their reduced adherence capacity. After this stabilization period, 10,000 low-passage cells (<10) were also seeded in 48-well plates to monitor and quantify their proliferation on days 3, 6, and 9.

### Cell Viability

Cell viability was determined by the trypan blue exclusion staining assay. Senescent cells were seeded according to the protocols described above. Senescent and proliferating cells were detached by treatment with 0.05% trypsin for 5 minutes; trypsin was neutralized with an equal proportion of medium with 10% FBS. We mixed 10 μl of cell suspension to count cells with 10 μl of trypan blue. Then, we used 10 μl of this mixture in the Bio-Rad TC20™ automated counter (catalog: 145-0101).

### Senescence-Associated β-Galactosidase (SA-β-gal) Activity

The SA-β-gal assay was prepared following the protocol previously reported in the literature (Debacq-Chainiaux et al., 2009). The assay was carried out for cells treated with 200 μM H_2_O_2_ at 72 hours, defining established senescence as a percentage greater than 60% of positive cells. In the case of high passage cells (> 65), the assay was performed 24 hours after seeding, taking established senescence as a percentage greater than 60% of senescent cells.

### Immunofluorescence of Senescence Markers and Caspase 3

Senescence induction was carried out according to the described protocols, using poly-L-lysine-free glass coverslips in 6-well plates. Cells were fixed with 4% paraformaldehyde (PFA) for 15 minutes at room temperature. Afterward, they were washed twice with PBS and incubated with 0.1% Triton X-100 for 45 minutes to permeabilize the cells. Subsequently, the cells were washed three times with PBS. Next, they were incubated with 0.1% Tween and 3% albumin for 90 minutes to block non-specific sites, followed by three washes with PBS. All antibodies (AB) were used to measure the senescent state at a 1:250 dilution. The selected markers were Lamin B1 (HAB8982, abcam), p21 (#6246, Santa Cruz), and GLB1 (ab220283, abcam).

The anti-active caspase-3 antibody (ab32042, Abcam) was used to evaluate the apoptosis caused by the drugs at a 1:100 concentration. The ABs were diluted in 0.1% PBS-Tween, and each sample was incubated with 50 μl of AB for 24 hours at 4°C with gentle agitation, followed by three washes with PBS; the secondary ABs were used at a 1:1000 dilution; the following ABs were used: Goat anti-Mouse IgG (H+L) Cross-Adsorbed Secondary Antibody, Alexa Fluor™ 594 (A11005, Invitrogen), Goat anti-Mouse IgG (H+L) Cross-Adsorbed Secondary Antibody, Alexa Fluor™ 488 (A11001), and Alexa Fluor® 594 AffiniPure™ Goat Anti-Mouse IgG (H+L) (115-585-003, Jackson Immunoresearch) and Goat anti-Rabbit IgG (H+L) Cross-Adsorbed Secondary Antibody, Alexa Fluor™ 488 (A11008, Invitrogen). The secondary ABs were diluted in 0.1% PBS-Tween, and each sample was incubated with 50 μl for 1 hour at room temperature in complete darkness and without agitation, followed by three washes with PBS. The coverslips with the cells were mounted on 50 μl of the mounting solution on glass slides. The mounting solution was prepared using 500 μl of distilled water and 500 μl of glycerol, to which 1 μl of 4′,6-diamidino-2-phenylindole (DAPI) (D1306, Invitrogen) was added. The coverslips with the cells were mounted on 50 μl of the mounting solution on glass slides.

Images were obtained using the Zeiss AXIOVERT 40 CFL microscope and processed with the *ZEN 2.3 (Blue edition)* software. A Python script was performed for fluorescence quantification and statistical analysis.

### Drug Preparation

For the biological assays, the following compounds were used: Fluorouracil (PHR1227, Sigma Aldrich), PF-562,271 (PZ0387, Sigma Aldrich), Dasatinib (CDS023389, Sigma Aldrich), and Quercetin (PHR1488, Sigma Aldrich). All compounds to be evaluated were dissolved in 100% dimethyl sulfoxide (DMSO) (D5879, Sigma Aldrich) to obtain a 100 mM stock solution. Subsequently, dilutions of each drug were prepared in 10% FBS culture medium without antibiotics, reaching a final concentration of 100 μM with 0.1% DMSO. For the positive control, a combination of Dasatinib and Quercetin (D+Q) with concentrations of 250 nM for D and 50 μM for Q was used (Moaddel et al., 2022). Additionally, medium with FBS without antibiotics and 0.1% DMSO was prepared. All solutions were filtered with a 0.22 μm pore size filter. The different concentrations used to evaluate senolytic activity were obtained by serial dilutions from the 100 μM solution.

### MTT Assay

Approximately 15,000 cells were seeded in 96-well plates for both SIPS (day 6) and RS and approximately 10,000 proliferating cells (passage < 10) were seeded. The RS and proliferating cells were incubated for 24 hours at 37°C and 5% CO_2_ before treatment application. After this period, the proliferating cells reached approximately a population of 15,000 cells. 200 μl of each drug for each concentration studied, and the vehicle group (0.1% DMSO) and the positive control (D+Q) were added. Based on a standard protocol for evaluating senolytic compounds, the cells were incubated for 48 hours under the same temperature and CO_2_ conditions (Yousefzadeh et al., 2018). After this time, a double wash with PBS was performed. A 3-(4,5-dimethylthiazol-2-yl)-2,5-diphenyltetrazolium bromide (MTT) (M2128, Invitrogen) solution was prepared at a final concentration of 0.5 mg/mL and pH 7.4, which was passed through a 0.22 μm filter. 100 μl of the MTT solution was added to each well, and the cells were incubated for 2 hours at 37°C and 5% CO_2_ in complete darkness. After incubation, the MTT was removed, and 150 μl of DMSO was added. The plates were gently shaken for 10 minutes and then incubated for another 15 minutes at 37°C. The relative optical density was quantified using a Multiskan SkyHigh microplate spectrophotometer (catalog number: A51119500C, Thermo Scientific). Before reading the absorbance at 570 nm, the plates were shaken for 10 seconds. Each drug at each concentration was evaluated in three independent experiments in triplicate.

The spectrophotometer readings were processed as follows: the Shapiro-Wilk normality test was applied, considering a *p*-value < 0.05 as significant. Then, a one-way ANOVA test was performed, accepting a *p*-value < 0.05 as significant. This was followed by Tukey’s multiple comparisons test with a Benjamini-Hochberg (FDR) adjustment to reduce the false positive rate, also considering a *p*-value < 0.05 as significant. All analyses were performed using Prism 8.0.2

## Results

### Identification of New SCAPs Candidates Through Transcriptomic Signature Profile Analysis

We based our approach on the hypothesis that silencing genes that promote the selective death of SCs generates a similar transcriptomic response to identify new SCAPs. To verify our hypothesis, we employed transcriptomic signature data obtained from the LINCS L1000 project. We used the consensus signature profiles, which, through robust z-scores, condensed the signature profiles of 9 cell lines into a single profile.

The first step was to determining similarities between the signatures caused by inhibiting known SCAPs genes. As previously mentioned, we employed the distance correlation (dCor) to measure this similarity, a measure capable of detecting both linear and non-linear associations. In Figure 2A, we present a hierarchical clustering analysis based on the median dCor, which revealed the existence of two clusters: the first consists of 15 genes (BCL2, GK3B, ABL1, CHEK1, PIK3CB, FYN, HIF1A, KIT, EFNB3, PIK3CG, AKT1, PTK2, ABL2, KDR, and SRC) with a median dCor = 0.390, *p* = 3×10-5, and the second group includes 6 genes (PIK3CA, PPM1L, FLT1, SERPINB2, BCL2L11, and MCL1) with a median dCor = 0.469, *p* = 0.00073. Considering both clusters, the 21 genes have a median dCor = 0.305, which is higher than expected by chance, *p* = 0.00054. The clustering based on the signature profiles of the SCAPs genes suggests the existence of multiple independent survival and anti-apoptotic resistance mechanisms; however, some genes showed high similarities in their signature patterns. Therefore, we decided to group the two identified clusters into a single group of 21 genes.

**Figure 2.**
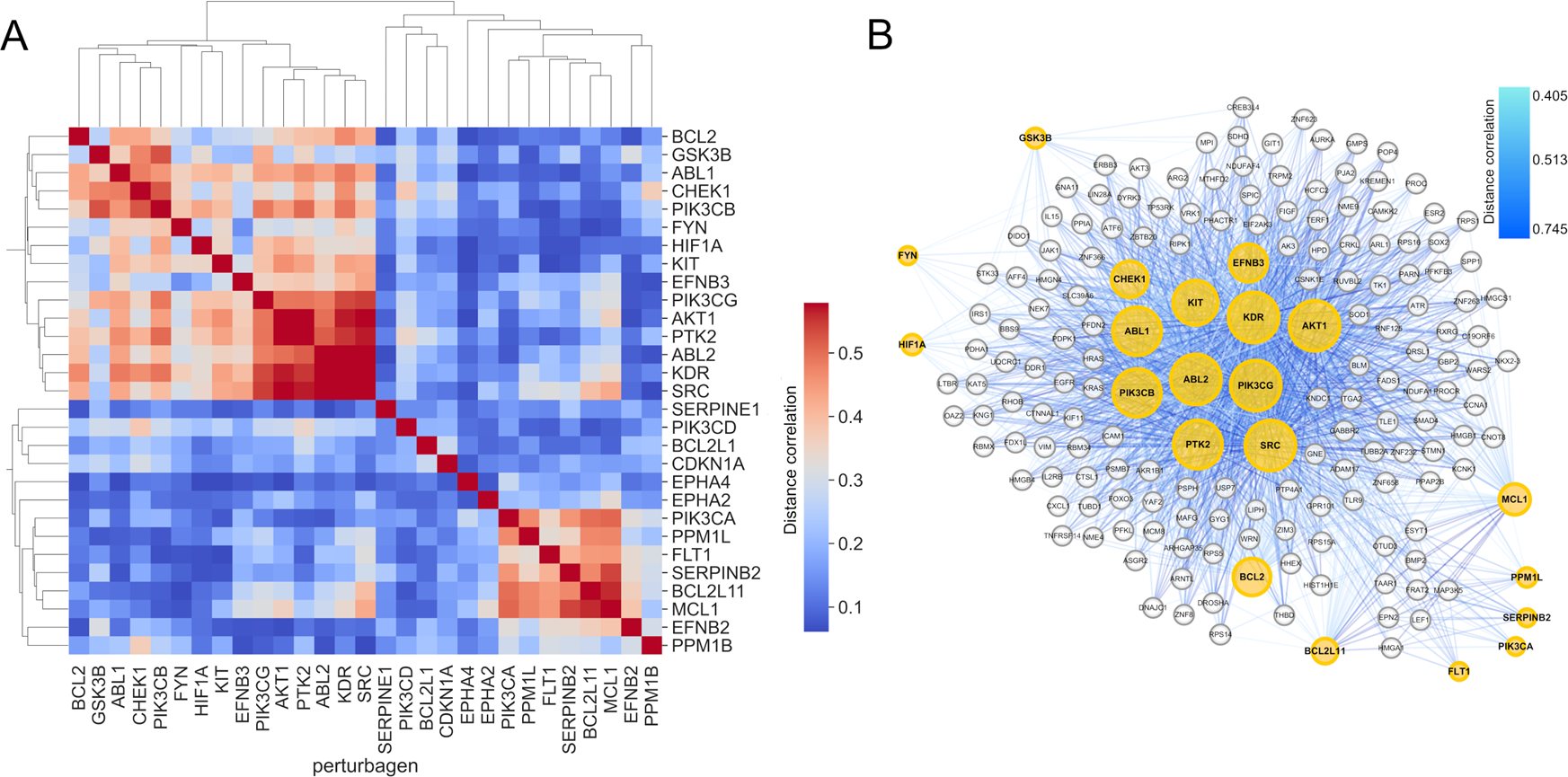
Similarity of the transcriptomic signatures of SCAPs. **A.** Hierarchical clustering analysis based on correlation distance. A group of 15 similar SCAPs genes was identified, showing a median dCor = 0.390, *p* = 3×10^-5^, and another group of 6 SCAPs genes with a median dCor = 0.469, *p* = 0.00073. The 21 SCAPs genes present a median dCor = 0.305, *p* = 0.00054. **B**. The inhibition of 154 individual genes (gray nodes) causes a transcriptomic signature similar (dCor > 0.405) to that observed when inhibiting the 21 SCAPs represented by yellow nodes, where the size of these nodes indicates the number of individual genes showing a similar signature profile.

Having signature profiles generated by the inhibition of 4,462 individual genes, we can identify those genes that, when inhibited, cause a signature similar to that observed in the group of 21 original SCAPs genes, which could trigger the same cellular response, selective death of SCs. We then calculated the median similarity in the signature of each of the 4,462 genes concerning the group of 21 SCAPs. Figure 2B shows the 154 genes (the complete list is in the supplementary material S2) that, when inhibited, generate a transcriptomic response like the inhibition of the 21 original SCAPs genes together, showing a dCor > 0.405 (cutoff established in materials and methods). The similarity between these genes is visualized through a network, where the size of the yellow nodes (SCAPs) reflects the number of genes (gray nodes) that induce a signature similar to that of the SCAPs genes.

### Gene Co-expression Network (GCN) Analysis

The individual inhibition of the 39 SCAPs genes triggers the selective death of SCs and causes a similar transcriptomic signature. These findings suggest a functional relationship among the SCAPs genes. For this reason, we decided to explore the correlation between these genes. For this purpose, we constructed gene co-expression networks (GCNs) using gene expression data from lung tissue of young (20 - 39 years) and old (60 - 70 years) individuals obtained from the GTEx project. We used mutual information (MI) as a metric to quantify the statistical dependence between gene pairs, as shown in Figure 3. Panel A shows the GCNs for young and old individuals. Interestingly, the SCAPs genes are co-expressed in both age groups; however, the network of old individuals loses connections, which could suggest that the evasion of programmed cell death and survival of SCs is an emerging property in response to changes in the functional coordination of these genes.

**Figure 3.**
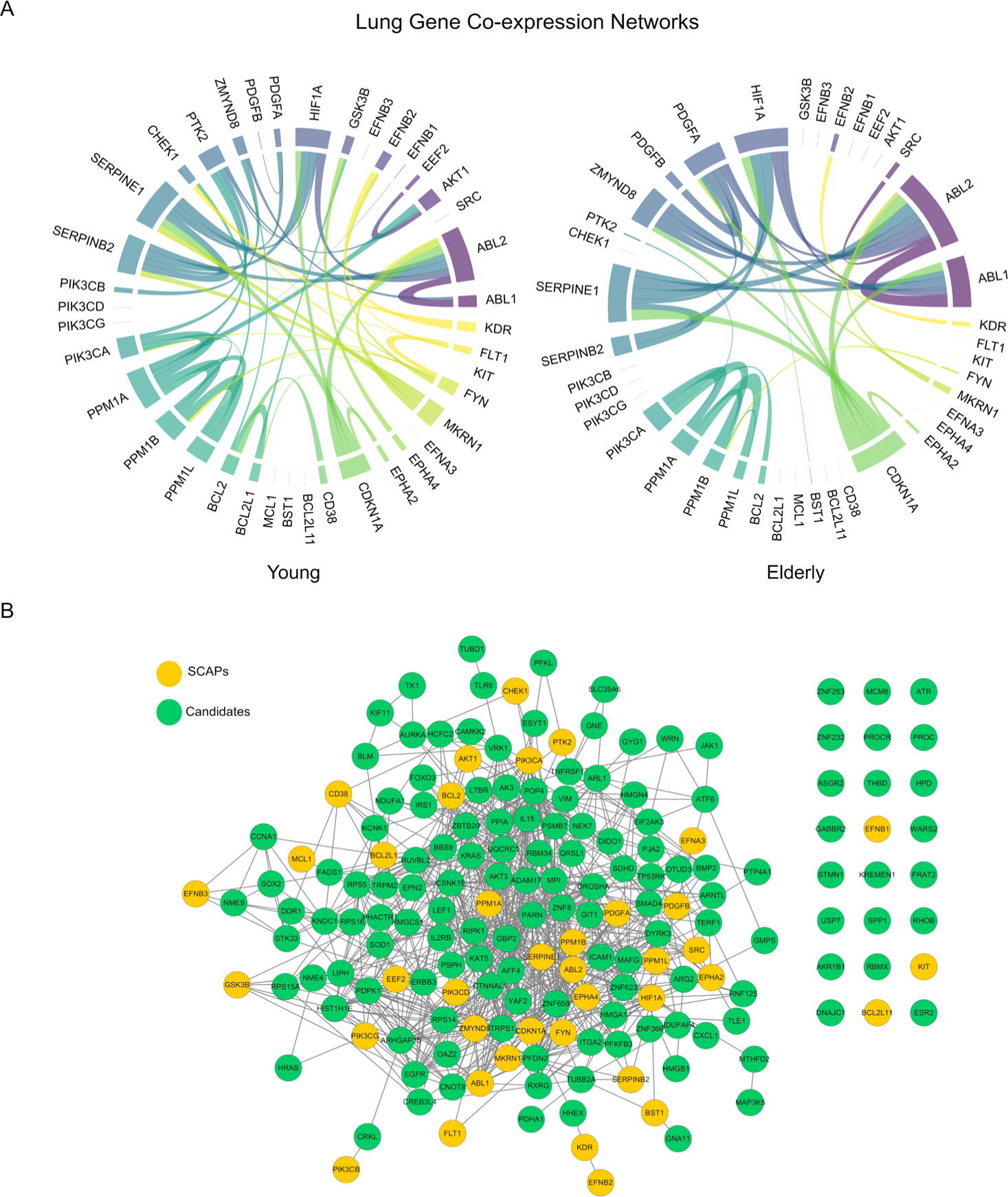
Gene Co-expression Network (GCN) Analysis. **A**. GCNs of the 39 SCAPs genes in human lung tissue from young (20-39 years) and elderly (60-70 years) individuals. The networks change their connection patterns with age, with the network of elderly individuals losing connections. **B**. GCN of genes from the LINCS L1000 dataset that, when inhibited, generate a transcriptomic signature comparable to that of SCAPs. Of the 154 genes, 120 are co-expressed with each other and with the original SCAPs genes, forming a dense network.

Based on this hypothesis, we investigated whether the 154 genes that induce a transcriptomic signature similar to that of the SCAPs genes are co-expressed with each other and with the SCAPs genes in the lung tissue of elderly individuals, assuming that there is a higher amount of SCs during this period of life and that apoptosis evasion is a consequence of the rewiring of GCNs. As shown in panel B, surprisingly, 120 of the 154 genes form part of a densely connected GCN. This result suggests that these 120 genes could be functionally related to the SCAPs genes and together form part of a transcriptional program potentially responsible for the anti-apoptotic capacity of SCs, so the inhibition of any of these 120 genes could lead to the selective death of SCs.

### Network Pharmacology

Identifying 120 potential SCAPs genes represents a major advance for discovering and repositioning senolytic molecules. As shown in Figure 4, we adopted a network pharmacology approach to explore this possibility.

**Figure 4.**
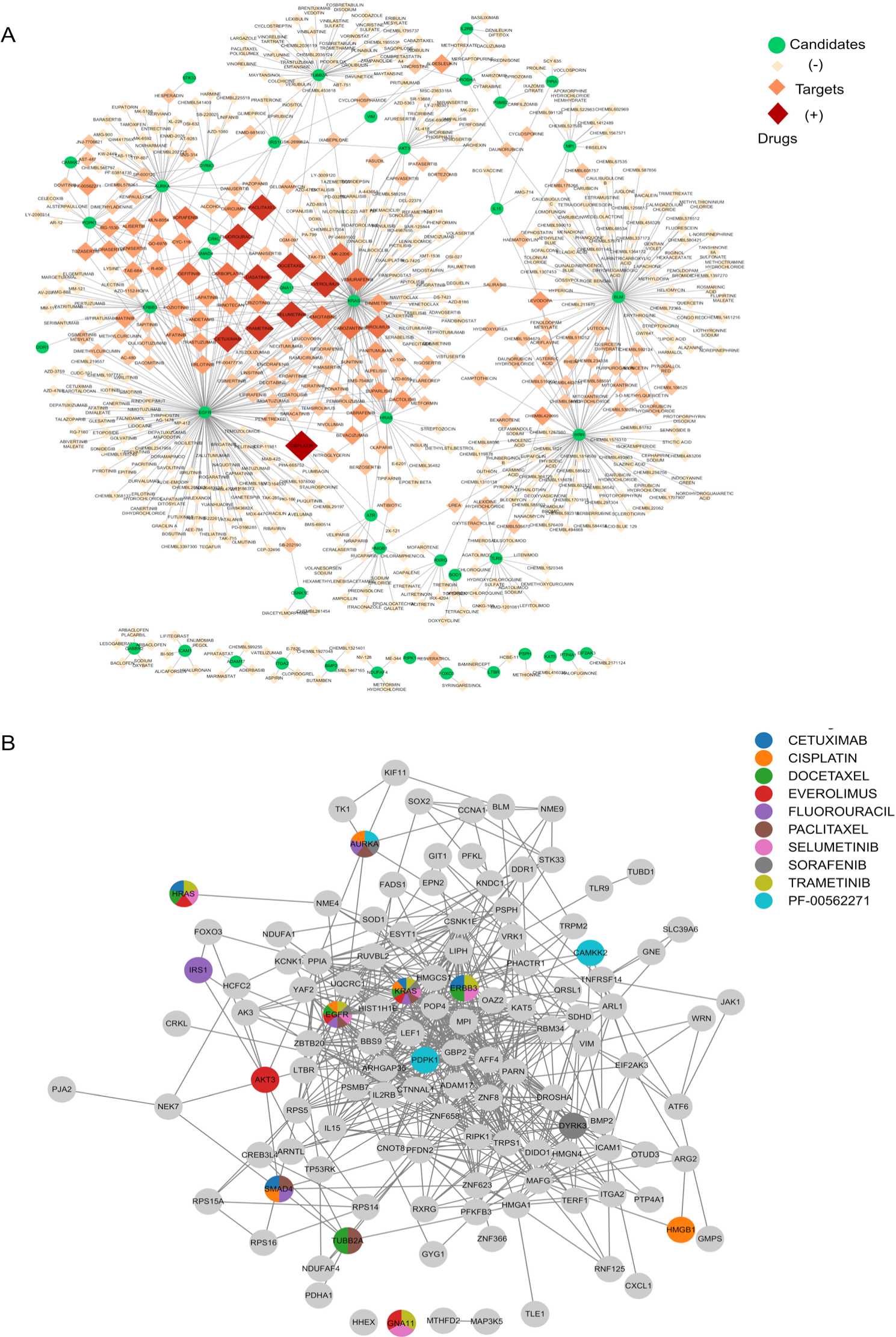
Drug-Target Networks **A**. A total of 620 molecules (red nodes) interacts with 44 of the 120 candidate genes (green nodes). Several of these molecules interact with more than one target, as observed through the nodes’ color and size gradient. **B.** Druggable target network, showing the 10 drugs with the highest number of targets within the GCN. The colors of pie-charted nodes represent the drugs that influence on those genes.

To explore whether our 120 candidates can function as pharmacological targets and direct small molecules to eliminate SCs selectively, we analyzed if they are targets of known senolytics. As shown in Supplementary Figure S1, we constructed a drug-target network that includes 550 targets of 59 senolytic drugs reported in the literature. Notably, 18 of our candidates (green nodes: KRAS, EGFR, JAK1, BLM, HRAS, TUBB2A, HMGB1, WRN, SMAD4, ERBB3, AKT3, VIM, CRKL, DYRK3, RUVBL2, DROSHA, IL15, and PPIA) are effective targets of senolytics, with an overlap greater than expected by chance (*p* = 0.0043, bootstrapping). This result validates that our methodology has succeeded in identifying targets that, when pharmacologically inhibited, induce selective death of SCs. Therefore, these 18 candidates are proposed as new SCAPs genes.

The fact that multiple genes from the 120 identified as potential SCAPs are targets of known senolytics allowed us to identify and propose molecules that, by inhibiting these targets, could act as potential senolytics. Using the DGIdb database, we identified molecules that interact with the candidate genes, as shown in Figure 4A. We found 620 molecules (red nodes) interacting with 44 of the 120 candidate genes. Notably, several of these molecules interact with more than one candidate gene, observed in the network through a color gradient and a progressive increase in node size, reflecting the number of targets with which each molecule interacts. The complete list of the 620 molecules and their respective targets is available in the supplementary material (File S2).

Using the GCN from Figure 3B and focusing only on the genes not belonging to the original SCAPs, and the 10 drugs with the highest number of targets identified in network 4B, we constructed a druggable target network, as shown in Figure 4C. We found that these 10 multi-target drugs interact with 14 genes from the GCN. Our results suggest that the pharmacological inhibition of these 14 targets could induce selective death of senescent cells, implying a potential senolytic effect for the drugs identified in network 4B. To validate the anti-apoptotic activity of the proposed targets, and the molecules with potential senolytic activity, we selected the molecules Fluorouracil and PF-562,271 based on criteria of cost, commercial availability, and the number of targets they inhibit.

### Experimental Validation

To validate our computational findings, we developed two senescence models using primary cultures of human lung fibroblasts from the CCD8-Lu cell line, as shown in Figure 5. For the stress-induced premature senescence (SIPS) model, oxidative stress was induced with H_2_O_2_. As observed in panel A, different concentrations (100, 150, and 200 μM) of H_2_O_2_ were evaluated. The 200 μM concentration was selected for senescence induction since, as shown in panels B and C, the cells exhibited characteristic senescence features and markers at this concentration. For the replicative senescence (RS) model, cells at a high passage (> 60) were used, showing a reduction in proliferation, as shown in panel D. Additionally, as shown in panels E and F, at this passage, the cells exhibited classic senescence features and markers.

**Figure 5.**
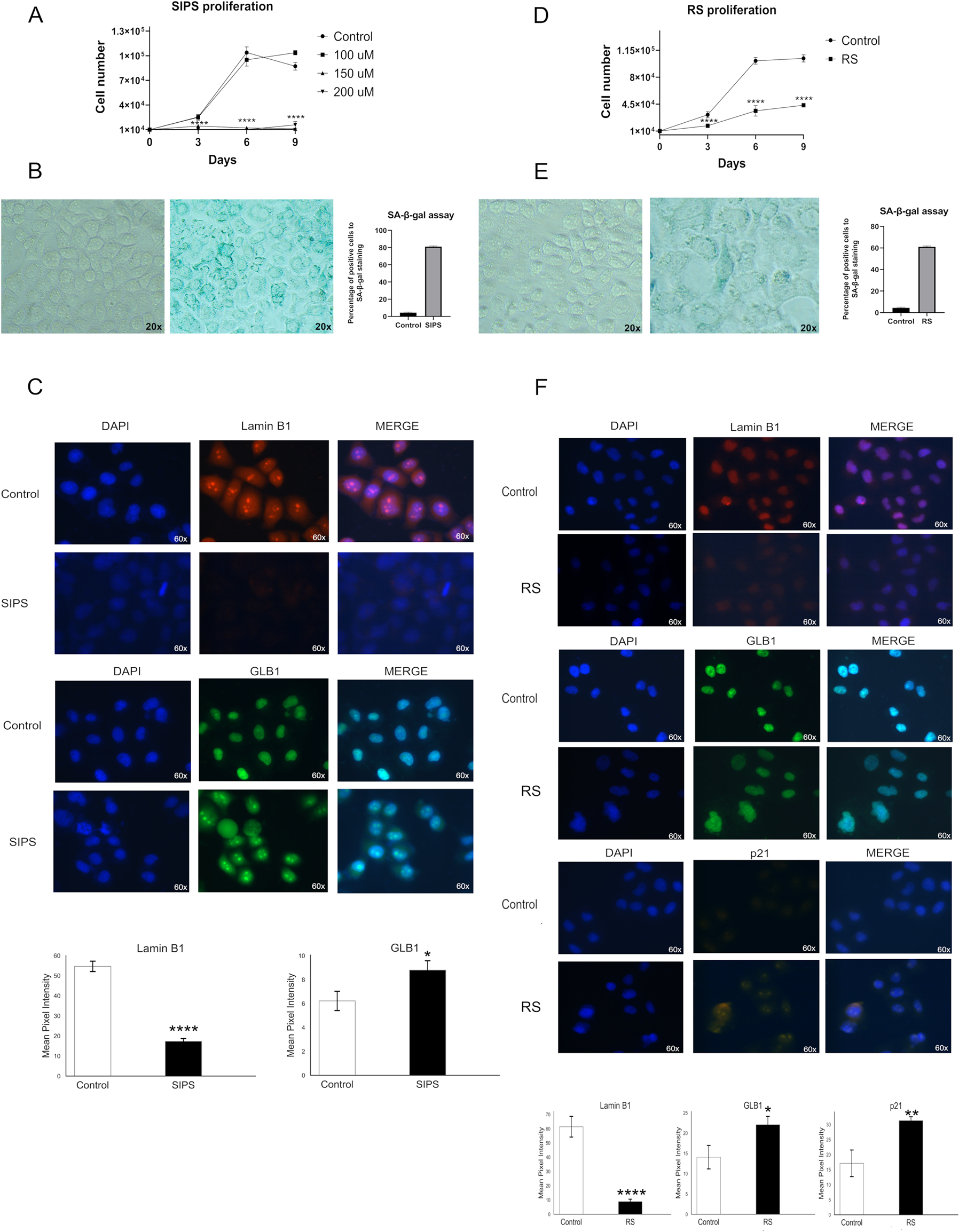
Standardization of two senescence models using the CCD8-Lu cell line. **A.** Cell proliferation curves with 100, 150, and 200 μM H_2_O_2_ treatments. A progressive reduction in proliferation is observed with increasing concentration. **B.** SA-β-gal assay of cells treated with 200 μM H_2_O_2_, around 80% of cells are positive, indicating a high incidence of senescence. **C.** Immunofluorescence of Lamin B1 and GLB1 markers in cells treated with 200 μM H2O2. **D.** Proliferation curves of cells at low passage (< 20) and high passage (> 60). **E.** SA-β-gal assay of cells treated with 200 μM H_2_O_2_, around 60% of cells are positive, confirming the effectiveness of the treatment in inducing senescence. **F.** Immunofluorescence of Lamin B1, GLB1, and p21 markers in cells treated with 200 μM H_2_O_2_, reaffirming the presence of senescent markers. The data are presented with the standard error of the mean (SEM) for n = 3 independent experiments performed in triplicate. Mann-Whitney U test. Statistical significance levels are indicated as *p* < 0.05, \**p* < 0.01, \*\*\**p* < 0.001, \*\*\*\**p* < 0.0001.

In the MTT assay, we evaluated the functionality of proliferating and senescent cells after being treated with three different concentrations of Fluorouracil (10, 12.5, and 15 μM) and PF-562,271 (1, 2.5, and 5 μM) for 48 hours. Furthermore, the D+Q combination was used as a control to compare senolytic effects, as shown in Figure 6. In panel A, the results indicate that neither Fluorouracil nor PF-562,271 exhibit a significant senolytic effect in the SIPS model; D+Q shows a slight effect, although it is not statistically significant (*p* = 0.067). However, as observed in panel B, treatment with 10 μM Fluorouracil for 48 hours significantly reduced the functionality of SCs (*p* < 0.0001) without impacting proliferating cells (*p* < 0.0001) in the RS model. The fact that Fluorouracil has a senolytic effect in RS but not SIPS suggests that the activation and development of SCAPs depend on the senescence-inducing stimulus. Interestingly, the senolytic effect was not intensified by increasing the concentration of Fluorouracil, although it decreased proliferating cells’ functionality.

**Figure 6.**
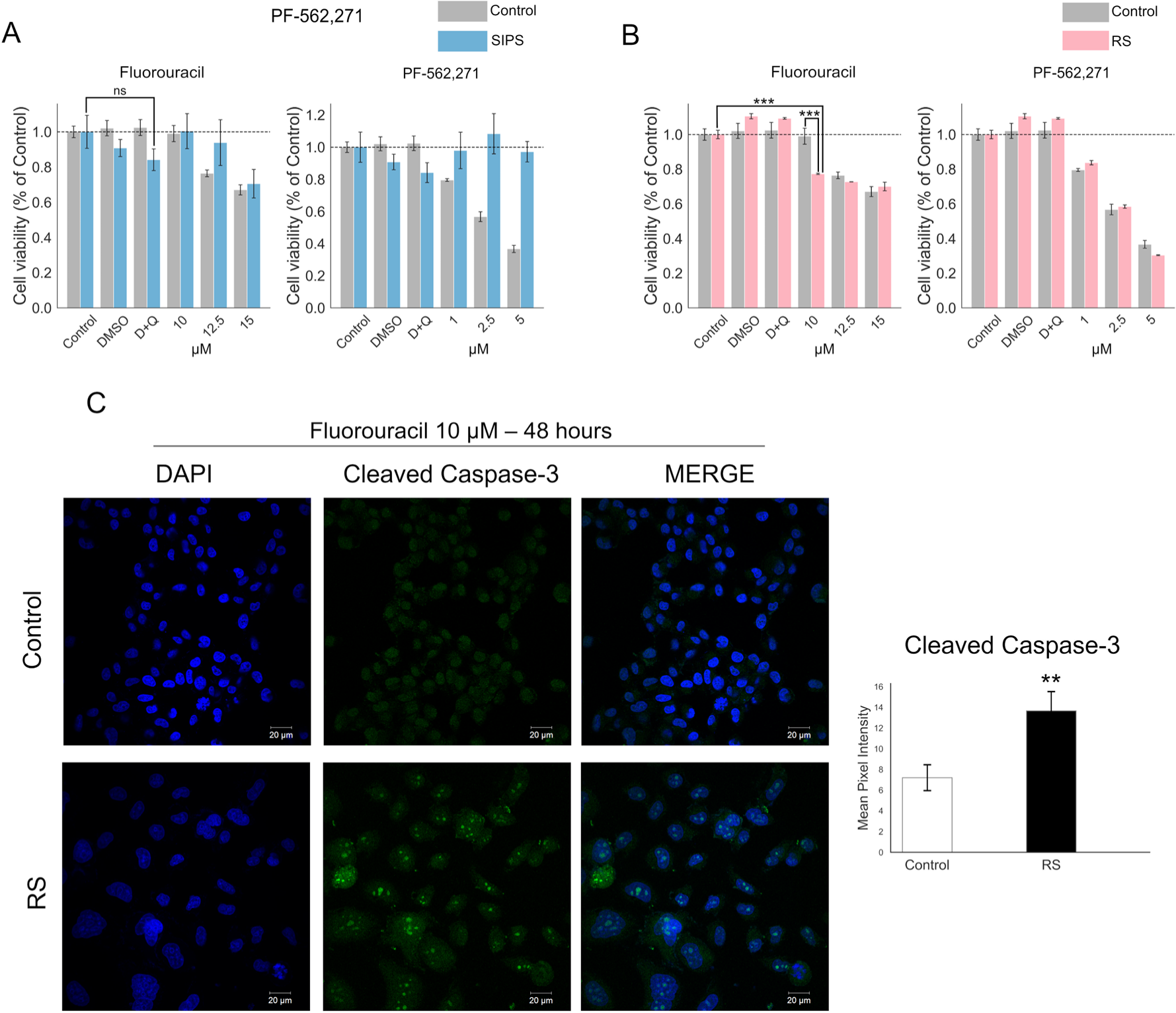
Effect of Fluorouracil and PF-562,271 at three different concentrations for 48 hours on the two senescence models. **A.** No effect of the two drugs was observed in the SIPS model. **B.** In the RS model, where Fluorouracil at a concentration of 10 μM significantly reduced cell functionality after 48 hours. **C.** Treatment with 10 μM Fluorouracil for 48 hours induced apoptosis in cells under RS but did not affect proliferating cells. The data are presented with the standard error of the mean (SEM) for n = 3 independent experiments performed in triplicate. Mann-Whitney U test. Statistical significance levels are indicated as *p* < 0.05, \**p* < 0.01, \*\*\**p* < 0.001.

On the other hand, PF-562,271 affected cells in the RS model, unlike the SIPS model; however, this effect was not selective, as it decreased the functionality of both proliferating and senescent cells. This result rules out its potential as a senolytic agent. In both models, it was observed that D+Q did not produce a senolytic effect.

As shown in panel C, we investigated whether the reduction in the functionality of SCs in the RS model after being treated with 10 μM Fluorouracil for 48 hours could be due to the selective induction of apoptosis. For this, we measured the expression of the active executioner Caspase-3, crucial in the apoptotic process, by immunofluorescence. The results showed that, in response to Fluorouracil treatment, the intensity of the active Caspase-3 signal in SCs increased 2.3-fold compared to non-senescent cells (*p* = 0.0093). This finding suggests that treatment with 10 μM Fluorouracil for 48 hours has a senolytic effect by inducing selective death of SCs; this points to the targets of this drug, including AURKA, EGFR, IRS1, SMAD4, KRAS, as potential new SCAPs.

## Discussion

Cellular senescence is a fundamental biological process that serves as a potent barrier against the proliferation of damaged cells, contributing significantly to aging and the suppression of tumorigenesis. Within this context, apoptosis resistance in SCs has garnered significant attention. This resistance is suggested to arise not from inherent cellular traits but rather from the aberrant behavior of certain genes known as SCAPs. This concept is supported by studies, which suggest that SCAPs play a crucial role in modulating cell survival mechanisms that enable senescent cells to evade programmed cell death (Rochette & Brash, 2008; Deryabin et al., 2021)

Further investigations indicate that the upregulation of specific survival and anti-apoptotic resistance genes and proteins can elucidate the mechanisms through which senescent cells successfully circumvent apoptosis (Hu et al., 2022; Roger et al., 2021; Chaib et al., 2022). This resistance to apoptosis is a key factor in understanding both the beneficial and detrimental roles of senescent cells in organisms. On the one hand, it prevents damaged cells from turning malignant, while on the other, it can contribute to age-related pathologies by allowing senescent cells to accumulate.Our results add to this understanding by suggesting that the development and activation of SCAPs are highly dynamic and involve complex networks of genes and proteins. These interactions, as evidenced in our analyses in Figures 2A and 3A, suggest that multiple genes and proteins work together to confer survival advantages to SCs. To further decipher how these genes promote apoptosis resistance, examining the specific molecular and cellular interactions that allow SCAPs to bypass programmed cell death effectively is crucial.

Studying SCAPs is therefore essential for elucidating the fundamental aspects of cellular aging and identifying potential therapeutic targets that can modulate senescence in age-related diseases and cancer. By understanding the intricate networks and pathways that underpin SCAPs function, researchers can develop strategies to enhance or inhibit these pathways, potentially leading to interventions that can mitigate the adverse effects of senescent cell accumulation or exploit the tumor-suppressive properties of cellular senescence.

Based on the results of the GCN analysis in Figure 3, we propose that the anti-apoptotic resistance of SCs is an emergent property derived from changes in co-expression patterns. These changes can alter existing patterns of functional association during the establishment and progression of senescence and may even lead to new patterns of functional association formation. In this way, we propose that the anti-apoptotic resistance of SCs is not an aberration but rather a well-defined, complexly regulated, and highly dynamic cellular program that could vary depending on the cell type, the senescence-inducing stimulus, and the stage of senescence, as has been reported with SASP (Maciel-Barón et al., 2016; Kumari & Jat, 2021). Moreover, considering the impact of the microenvironment on cellular senescence, a deeper analysis of how environmental factors contribute to the senescence and survival of senescent cells, particularly in age-related diseases and cancers, would enhance our understanding of therapeutic targets. Exploring how SCAPs genes interact with other cellular pathways critical for metabolism, differentiation, and inflammation could provide insights into their broader biological roles.

Considering the abovementioned concepts, we used various computational tools to discover new SCAPs. These approaches allowed us to identify 154 genes that, when individually inhibited with shRNA, trigger a transcriptomic response similar to that induced by silencing the original SCAPs genes. Of these genes, 120 are co-expressed with each other and with the SCAPs, which is relevant since genes that share co-expression patterns tend to be functionally related or regulated by the same transcriptional program (van Dam et al., 2018).

Of these 120 genes, we found that 18 are targets of known senolytics, which leads us to propose them as potential SCAPs. The discovery of these new SCAPs paves the way for the design and synthesis of new molecules and the identification of molecules in the experimental phase or drugs approved by regulatory agencies to reposition them as new senolytics. We have implemented a network pharmacology approach, which has revealed the existence of more than 600 novel molecules with potential senolytic activity. Many of these molecules can potentially act through a multi-target mechanism, a pharmacological paradigm that is particularly promising for combating complex diseases and phenotypes (Ramsay et al., 2018). This discovery prompts a closer examination of how these molecules could be integrated into existing or new treatment regimens for diseases associated with aging and cancer, enhancing their clinical and therapeutic relevance.

To demonstrate our computational findings’ validity and potential application, we conducted a proof-of-concept in two distinct senescence models, SIPS and RS, using two drugs identified in our network, Fluorouracil and PF-562,271. These experiments allowed us to explore and support at least three of our hypotheses: 1) The drugs from our network are potential senolytics. Treatment with 10 μM Fluorouracil for 48 hours reduced the functionality of cells in RS but did not affect proliferating cells; this reduction could be a consequence of the selective induction of apoptosis, as shown in Figure 5C2). Our candidates are new SCAPs: The senolytic effect of Fluorouracil is mediated through multi-target inhibition of AURKA, EGFR, IRS1, SMAD4, and KRAS. Interestingly, three of these proteins are effective targets of senolytics; AURKA is inhibited by the senolytic R-406 (Bamborough et al., 2008), EGFR is inhibited by the senolytics Geldanamycin (Puyo et al., 2008), MK-2206 (Parseghian et al., 2017), Ganetespib (Smith et al., 2015), and Curcumin (Wada et al., 2015). KRAS is inhibited by Panobinostat (Hayes et al., 2016), MK-2206 (Riquelme et al., 2016), Navitoclax (Horn et al., 2016), and Metformin (Iglesias et al., 2013). Although SMAD4 is not a target of any known senolytic, it has been shown to inhibit apoptosis (Du et al., 2020), as has IRS1, which, when inhibited, induces selective apoptosis in cancer cells (Scopim-Ribeiro et al., 2016). Finally, 3) The development of SCAPs depends on the senescence-inducing stimulus—figures 5A and 5B show that the tested molecules had differential effects. Despite the same experimental conditions, Fluorouracil exhibited a senolytic impact on the RS but not the SIPS model. A similar effect was observed with D+Q, which slightly impacted the SIPS model, although it was not statistically significant, suggesting that SCAPs are activated differentially depending on the senescence-inducing stimulus.

Studying SCAPs and their intricate interactions within cellular networks offers a promising avenue for advancing our understanding of cellular aging and the development of age-related pathologies. Our findings underscore the complexity of the senescence-associated anti-apoptotic mechanisms and highlight the potential for beneficially targeting these pathways to modulate senescence. As we continue to unravel the multifaceted roles of SCAPs in cellular health and disease, integrating this knowledge into clinical strategies could lead to innovative treatments that delay the onset of age-related diseases and enhance the quality of life in aging populations. Future research should focus on translating these molecular insights into practical applications, ensuring that the potential of senolytics to treat or prevent age-related conditions is fully realized.

### Conclusion and Perspectives

In this work, we propose that the development and activation of SCAPs result from changes in co-expression patterns that occur in response to the induction and progression of senescence. Therefore, the emergence of these transcriptional survival programs could be independent of the expression levels of their genes. Thus, targeting small molecules to the SCAPs GCNs could be a good pharmacological approach that allows the discovery and repositioning of new molecules with senolytic activity. This targeting strategy is supported by our findings of the multifunctional roles of these genes in cellular networks that influence metabolic, differentiation, and inflammatory pathways, underlining their potential broad impact on cell physiology.

We found that Fluorouracil acts as a senolytic through a multi-target mechanism involving the inhibition of AURKA, EGFR, IRS1, SMAD4, and KRAS. We consider these targets as new SCAPs since their transcriptomic signature profile is similar to that of the original SCAPs, their genes are co-expressed with the genes of the original SCAPs, forming a dense network, and their pharmacological inhibition leads to a reduction in the functionality of senescent cells. This decrease in functionality is attributed to the induction of selective apoptosis. The interaction of these genes within cellular networks provides a new understanding of how SCAPs contribute to cell survival under stress conditions induced by aging and pathological states. However, considering the polypharmacological nature of Fluorouracil, the senolytic effect could be given by inhibiting its other targets. Furthermore, the differential effects observed between the SIPS and RS models suggest that the context and type of senescence induction critically influence the effectiveness of senolytic interventions.

In our future studies, we aim to perform gene silencing experiments targeting AURKA, EGFR, IRS1, SMAD4, and KRAS using siRNA or shRNA across various cell types and in response to different stimuli that induce senescence. These experiments will deepen our understanding of their roles as SCAPs. Furthermore, we plan to employ advanced techniques like single cell sequencing alongside functional and comparative genomics approaches to gain more comprehensive insights into the dynamics and complexity of how survival and anti-apoptotic resistance networks develop. Through these approaches, we aim to elucidate the specific contributions of each SCAPs to the cellular resistance mechanisms and their potential as therapeutic targets. By integrating these methods, we will enhance our understanding of the components and mechanisms underpinning senescence and resistance to apoptosis.

## Supporting information

Proliferation

MTT

## Author Contributions

S.O.: Conceptualization, computational analyses, experimental validation, figure design, and article writing. M.K., J.P.V, N.E.L.D: Project direction and advisement, funding and material resources, writing, editing, review, and article approval. J.E.E. and H.T.: Advisement on computational analyses and article review. F.M.M: Material resources and article review. All authors approved the final version of the manuscript.

## Acknowledgments

We thank the Laboratorio de Supercómputo y Visualización en Paralelo (LSVP) of the Universidad Autónoma Metropolitana Campus Iztapalapa for the computing time granted on the ‘Yoltla’ supercomputer.

## Data availability

All data generated or analyzed during this study are included in the supplementary files of this published article.

## Code availability

The code used to support the analysis of this study can be found at https://github.com/Olascoaga/SCAPs

## References

Aguet, F., Barbeira, A. N., Bonazzola, R., Brown, A., Castel, S. E., Jo, B., Kasela, S., Kim-Hellmuth, S., Liang, Y., Oliva, M., Flynn, E. D., Parsana, P., Fresard, L., Gamazon, E. R., Hamel, A. R., He, Y., Hormozdiari, F., Mohammadi, P., Muñoz-Aguirre, M., … Volpi, S. (2020). The GTEx Consortium atlas of genetic regulatory effects across human tissues. Science, 369(6509), 1318–1330. 10.1126/SCIENCE.AAZ1776/SUPPL_FILE/AAZ1776_TABLESS10-S16.XLSX

Andonegui-Elguera, S. D., Zamora-Fuentes, J. M., Espinal-Enríquez, J., & Hernández-Lemus, E. (2021). Loss of Long Distance Co-Expression in Lung Cancer. Frontiers in Genetics, 12, 625741. 10.3389/FGENE.2021.625741/BIBTEX

Bamborough, P., Drewry, D., Harper, G., Smith, G. K., & Schneider, K. (2008). Assessment of chemical coverage of kinome space and its implications for kinase drug discovery. Journal of Medicinal Chemistry, 51(24), 7898–7914. 10.1021/JM8011036

Chaib, S., Tchkonia, T., & Kirkland, J. L. (2022). Cellular senescence and senolytics: the path to the clinic. Nature Medicine 2022 28:8, 28(8), 1556–1568. 10.1038/s41591-022-01923-y

Childs, B. G., Durik, M., Baker, D. J., & Van Deursen, J. M. (2015). Cellular senescence in aging and age-related disease: from mechanisms to therapy. Nature Medicine 2015 21:12, 21(12), 1424–1435. 10.1038/nm.4000

Coppé, J. P., Desprez, P. Y., Krtolica, A., & Campisi, J. (2010). The Senescence-Associated Secretory Phenotype: The Dark Side of Tumor Suppression. 10.1146/Annurev-Pathol-121808-102144, 5, 99–118. https://doi.org/10.1146/ANNUREV-PATHOL-121808-102144

Daniel Himmelstein, C. C. (n.d.). Computing consensus transcriptional profiles for LINCS L1000 perturbations. Thinklab. 10.15363/THINKLAB.D43

Debacq-Chainiaux, F., Erusalimsky, J. D., Campisi, J., & Toussaint, O. (2009). Protocols to detect senescence-associated beta-galactosidase (SA-βgal) activity, a biomarker of senescent cells in culture and in vivo. Nature Protocols 2009 4:12, 4(12), 1798–1806. 10.1038/nprot.2009.191

Deryabin, P. I., Shatrova, A. N., & Borodkina, A. V. (2021). Apoptosis resistance of senescent cells is an intrinsic barrier for senolysis induced by cardiac glycosides. Cellular and Molecular Life Sciences, 78(23), 7757. 10.1007/S00018-021-03980-X

Du, X., Li, Q., Yang, L., Liu, L., Cao, Q., & Li, Q. (2020). SMAD4 activates Wnt signaling pathway to inhibit granulosa cell apoptosis. Cell Death & Disease, 11(5). 10.1038/S41419-020-2578-X

Freshour, S. L., Kiwala, S., Cotto, K. C., Coffman, A. C., McMichael, J. F., Song, J. J., Griffith, M., Griffith, O. L., & Wagner, A. H. (2021). Integration of the Drug–Gene Interaction Database (DGIdb 4.0) with open crowdsource efforts. Nucleic Acids Research, 49(D1), D1144–D1151. 10.1093/NAR/GKAA1084

García-Cortés, D., Hernández-Lemus, E., & Enríquez, J. E. (2022). Loss of long-range co-expression is a common trait in cancer. BioRxiv, 2022.10.27.513947. 10.1101/2022.10.27.513947

Hayes, T. K., Neel, N. F., Hu, C., Gautam, P., Chenard, M., Long, B., Aziz, M., Kassner, M., Bryant, K. L., Pierobon, M., Marayati, R., Kher, S., George, S. D., Xu, M., Wang-Gillam, A., Samatar, A. A., Maitra, A., Wennerberg, K., Petricoin, E. F., … Der, C. J. (2016). Long-Term ERK Inhibition in KRAS-Mutant Pancreatic Cancer Is Associated with MYC Degradation and Senescence-like Growth Suppression. Cancer Cell, 29(1), 75–89. 10.1016/J.CCELL.2015.11.011

Henderson, A. R. (2005). The bootstrap: A technique for data-driven statistics. Using computer-intensive analyses to explore experimental data. Clinica Chimica Acta, 359(1–2), 1–26. 10.1016/J.CCCN.2005.04.002

Herranz, N., & Gil, J. (2018). Mechanisms and functions of cellular senescence. The Journal of Clinical Investigation, 128(4), 1238–1246. 10.1172/JCI95148

Horn, T., Ferretti, S., Ebel, N., Tam, A., Ho, S., Harbinski, F., Farsidjani, A., Zubrowski, M., Sellers, W. R., Schlegel, R., Porter, D., Morris, E., Wuerthner, J., Jeay, S., Greshock, J., Halilovic, E., Garraway, L. A., Caponigro, G., & Lehár, J. (2016). High-Order Drug Combinations Are Required to Effectively Kill Colorectal Cancer Cells. Cancer Research, 76(23), 6950–6963. 10.1158/0008-5472.CAN-15-3425

Hu, L., Li, H., Zi, M., Li, W., Liu, J., Yang, Y., Zhou, D., Kong, Q. P., Zhang, Y., & He, Y. (2022). Why Senescent Cells Are Resistant to Apoptosis: An Insight for Senolytic Development. Frontiers in Cell and Developmental Biology, 10, 822816. 10.3389/FCELL.2022.822816/BIBTEX

Iglesias, D. A., Yates, M. S., Van Der Hoeven, D., Rodkey, T. L., Zhang, Q., Co, N. N., Burzawa, J., Chigurupati, S., Celestino, J., Bowser, J., Broaddus, R., Hancock, J. F., Schmandt, R., & Lu, K. H. (2013). Another surprise from Metformin: novel mechanism of action via K-Ras influences endometrial cancer response to therapy. Molecular Cancer Therapeutics, 12(12), 2847–2856. 10.1158/1535-7163.MCT-13-0439

Kumari, R., & Jat, P. (2021). Mechanisms of Cellular Senescence: Cell Cycle Arrest and Senescence Associated Secretory Phenotype. Frontiers in Cell and Developmental Biology, 9, 645593. 10.3389/FCELL.2021.645593/BIBTEX

Maciel-Barón, L. A., Morales-Rosales, S. L., Aquino-Cruz, A. A., Triana-Martínez, F., Galván-Arzate, S., Luna-López, A., González-Puertos, V. Y., López-Díazguerrero, N. E., Torres, C., & Königsberg, M. (2016). Senescence associated secretory phenotype profile from primary lung mice fibroblasts depends on the senescence induction stimuli. *Age (Dordrecht*, Netherlands), 38(1), 1–14. 10.1007/S11357-016-9886-1

Margolin, A. A., Nemenman, I., Basso, K., Wiggins, C., Stolovitzky, G., Favera, R. D., & Califano, A. (2006). ARACNE: An algorithm for the reconstruction of gene regulatory networks in a mammalian cellular context. BMC Bioinformatics, 7(SUPPL.1), 1–15. 10.1186/1471-2105-7-S1-S7/FIGURES/6

Moaddel, R., Rossi, M., Rodriguez, S., Munk, R., Khadeer, M., Abdelmohsen, K., Gorospe, M., & Ferrucci, L. (2022). Identification of gingerenone A as a novel senolytic compound. PLOS ONE, 17(3), e0266135. 10.1371/JOURNAL.PONE.0266135

Olascoaga-Del Angel, K. S., Gutierrez, H., Königsberg, M., Pérez-Villanueva, J., & López-Diazguerrero, N. E. (2022). Exploring the fuzzy border between senolytics and senomorphics with chemoinformatics and systems pharmacology. Biogerontology, 23(4), 453–471. 10.1007/S10522-022-09974-X

Parseghian, C. M., Parikh, N. U., Wu, J. Y., Jiang, Z. Q., Henderson, L., Tian, F., Pastor, B., Ychou, M., Raghav, K., Dasari, A., Fogelman, D. R., Katsiampoura, A. D., Menter, D. G., Wolff, R. A., Eng, C., Overman, M. J., Thierry, A. R., Gallick, G. E., & Kopetz, S. (2017). Dual Inhibition of EGFR and c-Src by Cetuximab and Dasatinib Combined with FOLFOX Chemotherapy in Patients with Metastatic Colorectal Cancer. Clinical Cancer Research : An Official Journal of the American Association for Cancer Research, 23(15), 4146–4154. 10.1158/1078-0432.CCR-16-3138

Puyo, S., Le Morvan, V., & Robert, J. (2008). Impact of EGFR gene polymorphisms on anticancer drug cytotoxicity in vitro. Molecular Diagnosis & Therapy, 12(4), 225–234. 10.1007/BF03256288

Ramos-Carreño, C., & Torrecilla, J. L. (2023). dcor: Distance correlation and energy statistics in Python. SoftwareX, 22, 101326. 10.1016/j.softx.2023.101326

Riquelme, E., Behrens, C., Lin, H. Y., Simon, G., Papadimitrakopoulou, V., Izzo, J., Moran, C., Kalhor, N., Jack Lee, J., Minna, J. D., & Wistuba, I. I. (2016). Modulation of EZH2 Expression by MEK-ERK or PI3K-AKT Signaling in Lung Cancer Is Dictated by Different KRAS Oncogene Mutations. Cancer Research, 76(3), 675–685. 10.1158/0008-5472.CAN-15-1141

Rochette, P. J., & Brash, D. E. (2008). Progressive apoptosis resistance prior to senescence and control by the anti-apoptotic protein BCL-xL. Mechanisms of Ageing and Development, 129(4), 207–214. 10.1016/J.MAD.2007.12.007

Roger, L., Tomas, F., Gire, V., Galderisi, U., & Bernardo, G. Di. (2021). Mechanisms and Regulation of Cellular Senescence. International Journal of Molecular Sciences 2021, Vol. 22, Page 13173, 22(23), 13173. 10.3390/IJMS222313173

Scopim-Ribeiro, R., Machado-Neto, J. A., Eide, C. A., Campos, P. de M., Scheucher, P. S., Saad, S. T. O., Rego, E. M., Tognon, C. E., Druker, B. J., & Traina, F. (2016). Pharmacological IRS1/2 Inhibition Induces Apoptosis in BCR-ABL1T315I mutant Cells. Blood, 128(22), 1886–1886. 10.1182/BLOOD.V128.22.1886.1886

Shannon, P., Markiel, A., Ozier, O., Baliga, N. S., Wang, J. T., Ramage, D., Amin, N., Schwikowski, B., & Ideker, T. (2003). Cytoscape: A Software Environment for Integrated Models of Biomolecular Interaction Networks. Genome Research, 13(11), 2498–2504. 10.1101/GR.1239303

Smith, D. L., Acquaviva, J., Sequeira, M., Jimenez, J. P., Zhang, C., Sang, J., Bates, R. C., & Proia, D. A. (2015). The HSP90 inhibitor ganetespib potentiates the antitumor activity of EGFR tyrosine kinase inhibition in mutant and wild-type non-small cell lung cancer. Targeted Oncology, 10(2), 235–245. 10.1007/S11523-014-0329-6

Soto-Gamez, A., Quax, W. J., & Demaria, M. (2019). Regulation of Survival Networks in Senescent Cells: From Mechanisms to Interventions. Journal of Molecular Biology, 431(15), 2629–2643. 10.1016/J.JMB.2019.05.036

Subramanian, A., Narayan, R., Corsello, S. M., Peck, D. D., Natoli, T. E., Lu, X., Gould, J., Davis, J. F., Tubelli, A. A., Asiedu, J. K., Lahr, D. L., Hirschman, J. E., Liu, Z., Donahue, M., Julian, B., Khan, M., Wadden, D., Smith, I. C., Lam, D., … Golub, T. R. (2017). A Next Generation Connectivity Map: L1000 Platform and the First 1,000,000 Profiles. Cell, 171(6), 1437–1452.e17. 10.1016/J.CELL.2017.10.049

Székely, G. J., Rizzo, M. L., & Bakirov, N. K. (2007). Measuring and testing dependence by correlation of distances. 10.1214/009053607000000505, 35(6), 2769–2794. https://doi.org/10.1214/009053607000000505

van Dam, S., Võsa, U., van der Graaf, A., Franke, L., & de Magalhães, J. P. (2018). Gene co-expression analysis for functional classification and gene-disease predictions. Briefings in Bioinformatics, 19(4), 575–592. 10.1093/BIB/BBW139

Wada, K., Lee, J. Y., Hung, H. Y., Shi, Q., Lin, L., Zhao, Y., Goto, M., Yang, P. C., Kuo, S. C., Chen, H. W., & Lee, K. H. (2015). Novel curcumin analogs to overcome EGFR-TKI lung adenocarcinoma drug resistance and reduce EGFR-TKI-induced GI adverse effects. Bioorganic & Medicinal Chemistry, 23(7), 1507–1514. 10.1016/J.BMC.2015.02.003

Waskom, M. L. (2021). seaborn: statistical data visualization. Journal of Open Source Software, 6(60), 3021. 10.21105/JOSS.03021

Wissler Gerdes,E. O., Zhu, Y., Weigand, B. M., Tripathi, U., Burns, T. C., Tchkonia, T., & Kirkland, J. L. (2020). Cellular senescence in aging and age-related diseases: Implications for neurodegenerative diseases. International Review of Neurobiology, 155, 203–234. 10.1016/BS.IRN.2020.03.019

Yousefzadeh, M. J., Zhu, Y., McGowan, S. J., Angelini, L., Fuhrmann-Stroissnigg, H., Xu, M., Ling, Y. Y., Melos, K. I., Pirtskhalava, T., Inman, C. L., McGuckian, C., Wade, E. A., Kato, J. I., Grassi, D., Wentworth, M., Burd, C. E., Arriaga, E. A., Ladiges, W. L., Tchkonia, T., … Niedernhofer, L. J. (2018). Fisetin is a senotherapeutic that extends health and lifespan. EBioMedicine, 36, 18–28. 10.1016/j.ebiom.2018.09.015

Zhu, Y., Tchkonia, T., Pirtskhalava, T., Gower, A. C., Ding, H., Giorgadze, N., Palmer, A. K., Ikeno, Y., Hubbard, G. B., Lenburg, M., O’hara, S. P., Larusso, N. F., Miller, J. D., Roos, C. M., Verzosa, G. C., Lebrasseur, N. K., Wren, J. D., Farr, J. N., Khosla, S., … Kirkland, J. L. (2015). The Achilles’ heel of senescent cells: from transcriptome to senolytic drugs. Aging Cell, 14(4), 644–658. 10.1111/ACEL.12344

