## Supplementary material for "Transcriptomic signatures and network-based methods uncover new Senescent Cell Anti-Apoptotic Pathways and Senolytics": S1

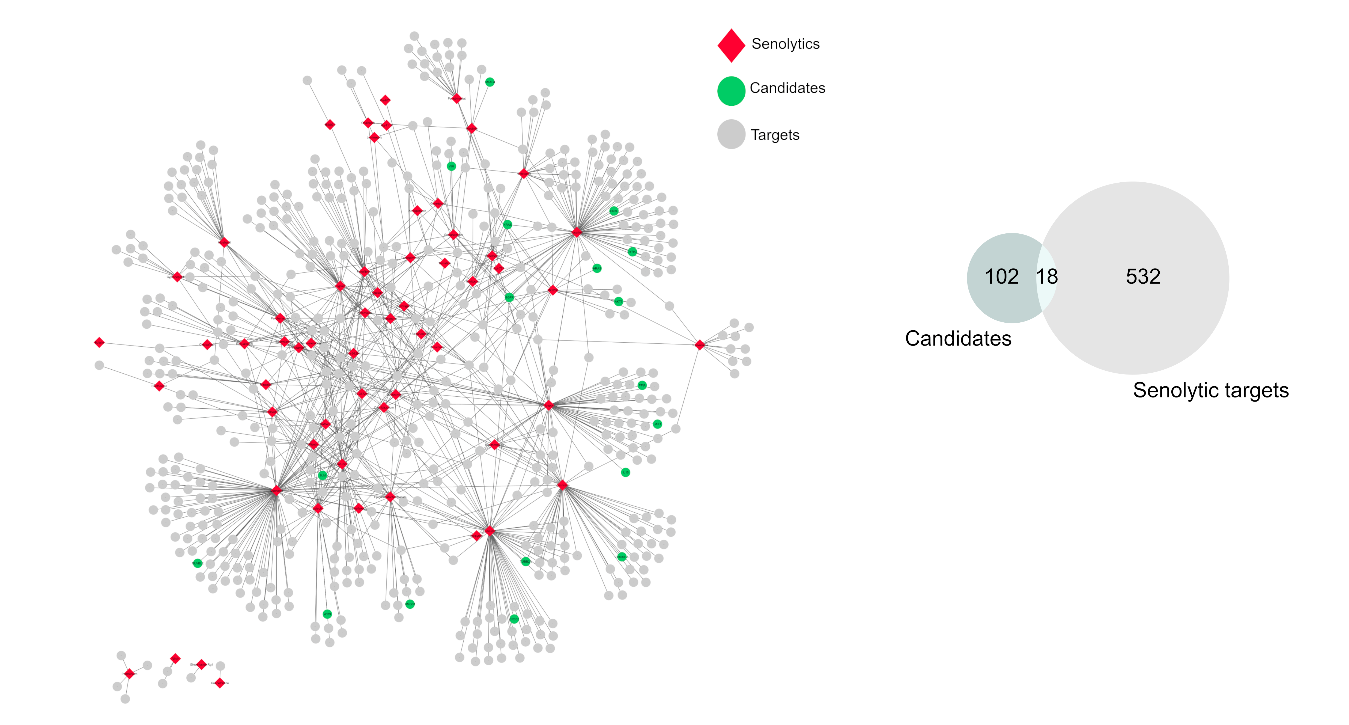

**Figure S1.** Drug – Target Network. network that includes 550 targets of 59 senolytic drugs reported in the literature. Notably, 18 of our candidates (green nodes: KRAS, EGFR, JAK1, BLM, HRAS, TUBB2A, HMGB1, WRN, SMAD4, ERBB3, AKT3, VIM, CRKL, DYRK3, RUVBL2, DROSHA, IL15, and PPIA) are effective targets of senolytics, with an overlap greater than expected by chance (*p = 0.0043*, bootstrapping)
